# Microclimates can be accurately predicted across ecologically important remote ecosystems

**DOI:** 10.1101/2021.01.14.426699

**Authors:** D.J. Baker, C.R. Dickson, D.M. Bergstrom, J. Whinam, I.M.D Maclean, M.A. McGeoch

## Abstract

Microclimate information is often crucial for understanding ecological patterns and processes, including under climate change, but is typically absent from ecological and biogeographic studies owing to difficulties in obtaining microclimate data. Recent advances in microclimate modelling, however, suggest that microclimate conditions can now be predicted anywhere at any time using hybrid physically- and empirically-based models. Here, for the first time, we test the utility of this approach across a remote, inaccessible, and climate change threatened polar island ecosystem at ecologically relevant scales. Microclimate predictions were generated at a 100 × 100 m grain (at a height of 4 cm) across the island, with models parameterised using either meteorological observations from the island’s weather station (AWS) or climate reanalysis data (CRA). AWS models had low error rates and were highly correlated with observed seasonal and daily temperatures (root mean squared error of predicted seasonal average T_mean_ ≤ 0.6 °C; Pearson’s correlation coefficient (r) for the daily T_mean_ ≥ 0.86). By comparison, CRA models had a slight warm bias in all seasons and a smaller diurnal range in the late summer period than *in situ* observations. Despite these differences, the modelled relationship between the percentage cover of the threatened endemic cushion plant *Azorella macquariensis* and microclimate varied little with the source of microclimate data (r = 0.97), suggesting that both model parameterisations capture similar patterns of spatial variation in microclimate conditions across the island ecosystem. Here, we have shown that the accurate prediction of microclimate conditions at ecologically relevant spatial and temporal scales is now possible using hybrid physically- and empirically-based models across even the most remote and climatically extreme environments. These advances will help add the microclimate dimension to ecological and biogeographic studies, which could be critical for delivering climate change-resilient conservation planning in climate-change exposed ecosystems.

## INTRODUCTION

Microclimates created by landscape and vegetation structures play a key role in facilitating the persistence of species in locations that would otherwise be climatically inhospitable (Maclean, Hopkins, Bennie, Lawson, & Wilson, 2015; Suggitt et al., 2018). Without information on microclimates, we observe only course-scale associations between biodiversity and climate, omitting important fine-scale conditions that directly affect organisms (Bütikofer et al., 2020; Storlie et al., 2014). Understanding the role of microclimates in structuring biodiversity and facilitating the persistence of species under climate change is vital (Storlie et al., 2014), but studies to date have been limited in geographical and temporal scope by an absence of microclimate data in most locations and for most time periods. However, recent advances in microclimate modelling suggest that microclimate conditions can now be predicted anywhere at any time (Lembrechts & Lenoir, 2020). This claim has profound consequences for biodiversity conservation because climate change is already impacting ecosystems across the globe, and the availability of spatially and temporally explicit microclimate information could help provide a crucial and timely advance in our understanding of how biodiversity is likely to respond.

Perhaps nowhere is this advance needed more urgently than in remote and climate change threatened polar ecosystems (Cahoon, Sullivan, Shaver, Welker, & Post, 2012; Chown et al., 2015; Niittynen et al., 2020). These isolated and climatically extreme environmental, notably the terrestrial ecosystems of the Southern Ocean, have long been regarded as sentinels of emerging climate change impacts (Bergstrom & Chown, 1999). With the polar regions having already experienced rapid warming (Clem et al., 2020; Turner et al.,2014), the impacts to biodiversity are now clearly evident (Bergstrom et al., 2015; Descamps et al., 2017; McClelland et al., 2018) and are predicted to increase in severity under future climate change scenarios (Duffy et al., 2017; Niskanen, Niittynen, Aalto, Väre, & Luoto, 2019; Wauchope et al., 2017). Temperature is the dominant factor shaping the distribution of species in these climatically extreme regions, and the effects are realised at the scales of both macroclimate (Leihy, Duffy, & Chown, 2018) and microclimate (Hilde et al., 2016; Kankaanpää, Abrego, Vesterinen, & Roslin, 2020; Niittynen et al., 2020; Nyakatya & McGeoch, 2008). Macroclimate effects are routinely studied, but owing to difficulties in obtaining microclimate data, the microclimate dimension is typically absent from analyses of species response to climate and predictions of changes in the spatial distribution and temporal dynamics of species across these extreme environments under climate change.

With sparse weather observations across remote and inaccessable regions, researchers have tended to use terrain proxies of microclimate (e.g. slope, aspect, elevation, and variables derived from these characteristics, such as wind shelter and topographic wetness index) for modelling species distributions (Bricher, Lucieer, Shaw, Terauds, & Bergstrom, 2013) or for evaluating the drivers of ecological processes (Dickson et al., 2019). Terrain proxies essentially assume stationarity in climate conditions during the period over which terrain is acting as a proxy; however, this is unlikely during a period of rapid climate change (Dobrowski, 2011). By comparison, the deployment of data loggers (e.g. measuring temperature and humidity) provide a more direct way to measure fine-scale climate conditions and have been used in particular to study the effects of fine-scale temperature and humidity conditions on focal species or ecosystems (Dickson et al., 2020; Niittynen et al., 2020; Nyakatya & McGeoch, 2008). However, data loggers provide information for point locations only and opportunities for deploying and maintaining microclimate sensor arrays in remote regions are extremely limited. Satellite remote sensing data by contrast provide extensive, contiguous spatial coverage of surface temperature conditions—more than 15 years of surface temperature observations measured at 1 to 2 day intervals—across polar regions (Leihy, Duffy, Nortje, & Chown, 2018), but are currently limited to a c. 1 km spatial resolution.

Hybrid physical- and empirical-based microclimate models offer a computationally efficient and theoretically grounded alternative for obtaining microclimate data across remote regions. These models use well-established physical relationships between near-ground temperatures and a reference air temperature (e.g. Maclean et al. 2019) and the modulation of these relationship by landscape physiography (Lembrechts & Lenoir, 2020), to accurately predict hourly temperatures at spatial grains one or two orders-of-magnitude lower than the best available satellite remote sensing data. These models, therefore, provide spatially contiguous fine scale estimates—even at a 1 m grain size (Kearney, Gillingham, Bramer, Duffy, & Maclean, 2020)—of biologically important microclimate variation at high temporal resolutions (i.e. hourly). Hybrid models use empirical data (e.g. local meteorological records and digital elevation) to derive parameter estimates for the physical equations that describe near-surface temperatures at any position in the landscape (Maclean, Suggitt, Wilson, Duffy, & Bennie, 2017). They predict the microclimate conditions that result from heat exchange at the microclimate scale, as well as mesoclimate effects—such as, elevation, cold-air drainage, and coastal effects—on the realised microclimate conditions (Maclean et al., 2017). Models are typically parameterised using *in situ* meteorological observations (e.g. networks of data loggers and meteorological stations), which permit accurate estimation of local conditions (Maclean, Mosedale, & Bennie, 2019). The reliance on *in situ* data has previously been a barrier to microclimate modelling in remote regions. However, with the availability of climate reanalysis data—comprehensive estimates of how weather and climate are changing over time derived from observations and numerical models—that can be used to parameterise microclimate models in these locations, microclimates can now be predicted in any location or time period covered by these data (Lembrechts & Lenoir, 2020).

Robust and comprehensive testing of novel methods are required before they can be used with confidence in ecological and biogeographic studies. Here, we test the ability of a hybrid physically- and empirically-based microclimate model to reconstruct hourly fine-scale climate conditions (c. 100 × 100 m at 4 cm above the surface) across a remote, inaccessible, and climate change threatened polar island ecosystem. The models are parameterised using:(1)meteorological data from the island’s automatic weather station (AWS) or (2) climate reanalysis data (CRA). We assess the ability of these two models, either with or without *in situ* meteorological observations, to predict microclimate conditions across a network of 62 *in situ* data loggers distributed across the entire island ecosystem. We demonstrate how differences in predictions from the two microclimate models can affect bioclimate variables and evaluate the consequences of these difference for predictions of the distribution of the threatened foundation species *Azorella macquariensis* Orchard (Apiaceae) across the island. We discuss the game-changing nature of these advances in microclimate modelling for ecological and biogeographical analysis, particularly in remote wilderness locations.

## METHODS

### Study system

Macquarie Island (158°55’E; 54°30’S) is situated in the Southern Ocean. The island covers an area of 128 km^2^, is c. 34 km long, c. 5 km wide, and has a maximum elevation of 433 m above sea level (asl; Chown et al., 1998). The island’s climate is cool, misty, and windy, with very low daily variations (Dickson et al., 2019; Selkirk, Seppelt, & Selkirk, 1990). Since 1970, the island’s climate has undergone significant changes, including increased precipitation, wind speed, and sunshine hours (Adams, 2009; Bergstrom et al., 2015).

### Meteorological Observations

The Australian Bureau of Meteorology station (#300004) is positioned at the northern isthmus of Macquarie Island (−54.4994 °S 158.9369 °W) at a height of 6 m asl. The thermometer and barometer are positioned at a height of 1.2 and 1.3 m above the ground, respectively. Manual records have been collected at the station since 1948 and an automated weather station (AWS) was installed in 1997. At hourly intervals, the AWS records air temperature, dew-point temperature, relative humidity, vapour pressure, wind speed and direction, station level pressure (later converted to pressure at sea level), and precipitation.

### Microclimate Model

Hourly temperatures were modelled at a 100 × 100 m resolution at a height of 4 cm above the ground using the *microclima* (Maclean et al., 2019) and *NicheMapR* (Kearney & Porter, 2017) R packages. In particular, we use the integration of these two packages within the *runauto* function of the *microclima* package (Kearney et al., 2020). This function uses temperatures estimated at each location by *NicheMapR* to calibrate the linear model relating the local temperature anomaly to the reference temperature as a function of net radiation and wind speed (Maclean et al., 2019), as calculated by *microclima*. Without this integration, the *microclima* microclimate model requires *in situ* mesoclimate and microclimate reference data, collected using an appropriate sampling strategy across the region of interest, to parameterise the model. In the fully automated model, National Centers for Environmental Prediction (NCEP; Kanamitsu et al., 2002; Kemp, Emiel van Loon, Shamoun-Baranes, & Bouten, 2012) climate reanalysis data is downscaled and interpolated to provide hourly information on the reference temperatures and atmospheric forcing conditions that are used to parameterise the model. NCEP (specifically NCEP-DOE reanalysis R-2 [hereafter, NCEP2]) derived variables include temperature at 2 m (°C), specific humidity at 2 m (Kg / Kg), surface pressure (Pa), wind speed at 2 m (m sec^-1^), wind direction (degrees from N), atmospheric emissivity, multiple measures of direct and diffuse radiation, and cloud cover (%). In our analysis, the ground and canopy albedo were fixed at 0.15 and 0.23 and habitat was specified as ‘Barren or sparsely vegetated’. The models were parameterised using either (Table 1a):

**Table 1.** Summary of datasets used in model parameterisation and the model evaluation statistics.

|  |  |
| --- | --- |
| a) | Microclimate model parameterisation data |
|  | **1) AWS (automatic weather station)** |
|  | AWS data + NCEP2* CRA data |
|  | **2) CRA (climate reanalysis)** |
|  | NCEP2 CRA data |
| b) | Model evaluation (62 sites) |
|  | **Seasonal statistics** |
| i. | Calculate seasonal mean of daily $T_{[\text{min}, \text{mean}, \text{max}]}$ for each site for observed vs. predicted and observed vs. macroclimate (i.e. NCEP2 derived temperatures). |
| ii. | Calculate the root mean squared error (RMSE) and median absolute error (MAE) for the observed vs. predicted and observed vs macroclimate across all 62 evaluation sites. |
|  | **Daily statistics** |
| i. | Calculate the RMSE and Pearson's correlation coefficient (r) between the observed vs. predicted and observed vs. macroclimate daily $T_{[\text{min}, \text{mean}, \text{max}]}$ for each site. |
| ii. | Calculate mean RMSE and r across all 62 evaluation sites. |

\* National Centers for Environmental Predictions(NCEP)-DOE reanalysis R-2

1. AWS – the hourly meteorological observations from the island’s AWS for air temperature, air pressure at sea level, wind direction, wind speed, and relative humidity (with hourly estimates of the emissivity of the atmosphere, radiation, and cloud cover downscaled from NCEP2 data).
2. CRA – hourly meteorological data obtained solely from downscaled NCEP2 data.

The NCEP2 data were downloaded and temporally interpolated from six-hourly to hourly values using the hourlyNCEP function from *microclima*. Models were run in hourly time steps for two three–month periods that define the island’s growing season: early summer (1^st^ October to 31^st^ December) and late summer (1^st^ January to 31^st^ March).

### Model Evaluation

Model predictions were evaluated against the *in situ* microclimate observation data (‘Obs’) across the 62 sites (see Table 1b for overview). A stratified random sampling approach was used to select the 62 microclimate monitoring sites across Macquarie Island’s plateau ecosystems, stratified by both terrain class and spatial blocks (Dickson et al., 2019). The stratification was designed to maximise the range of microclimate conditions sampled by the data loggers across combinations of latitude, elevation, and exposure (i.e. to moisture, wind, and solar radiation). We used DS1923 Hygrochon Temperature & Humidity iButtons (Maxim) that were set to record every 4 hours. The data-loggers were housed in light grey PVC jars and attached by a free hanging plastic fob. The jars had three small slats cut into each side to facilitate airflow, while still providing shelter from direct solar radiation and precipitation. The upturned jars and data-loggers were attached to stakes 4 cm above the ground, which is representative of the height of most of the plateau vascular and non-vascular flora. Microclimate data-loggers were deployed between 15/12/2016 and 27/02/2017 and collected between 24/11/2017 and 02/01/2018.

First, because seasonal averages are often used in ecological studies (e.g. Dickson et al., 2020), we assessed seasonal predictions by calculating the root mean squared error (RMSE) and median absolute error (MAE) of the seasonal average daily T_min_, T_mean_, and T_max_ between the observed and predicted and the observed and macroclimate (i.e. NCEP2 derived) temperatures across all 62 sites (Table 1b). Second, we assessed the predictive performance of the models at daily intervals across the complete time-series at each site using the RMSE and Pearson’s correlation coefficient (r) for each temperature variable (Table 1b); the mean and standard deviation of these within-site RMSE across the 62 sites (see below) are reported (Table 2). We also evaluated the spatial distribution of prediction errors for the observed and predicted temperature by mapping the observed minus predicted seasonal T_min_, T_mean_, and T_max_ for each season and model (i.e. AWS and CRA).

**Table 2.** Evaluation of the predictions from the microclimate model results against *in situ* microclimate data loggers. AWS = automated weather station; CRA = climate reanalysis; T_macro_ = macroclimate temperature (reference air temperature derived from NCEP2 time-series); Obs = *in situ* observations.

| | Metric | Season | AWS vs. Obs | | | CRA vs. Obs | | | Obs vs $T_{macro}$ | | |
| --- | --- | --- | --- | --- | --- | --- | --- | --- | --- | --- | --- |
| | | | $T_{min}$ | $T_{mean}$ | $T_{max}$ | $T_{min}$ | $T_{mean}$ | $T_{max}$ | $T_{min}$ | $T_{mean}$ | $T_{max}$ |
| Mean of seasonal statistics | RMSE (°C) | Early summer | 0.57 | 0.61 | 0.99 | 1.23 | 1.20 | 1.50 | 3.61 | 2.42 | 4.33 |
|  |  | Late summer | 0.27 | 0.45 | 0.77 | 0.59 | 0.27 | 0.82 | 2.80 | 1.47 | 3.33 |
|  | MAE (°C) | Early summer | 0.49 | 0.50 | 0.85 | 1.19 | 1.11 | 1.33 | 3.59 | 2.36 | 4.22 |
|  |  | Late summer | 0.22 | 0.38 | 0.59 | 0.54 | 0.22 | 0.62 | 2.78 | 1.40 | 3.21 |
| Mean across 62 sites of within-site daily statistics (± SD) | RMSE (°C) | Early summer | 1.13<br>(0.171) | 1.21<br>(0.259) | 2.23<br>(0.595) | 1.93<br>(0.138) | 1.67<br>(0.173) | 2.54<br>(0.473) | 3.30<br>(0.271) | 2.50<br>(0.334) | 3.09<br>(0.612) |
|  |  | Late summer | 0.77<br>(0.191) | 0.94<br>(0.221) | 2.23<br>(0.692) | 1.48<br>(0.171) | 1.23<br>(0.165) | 2.42<br>(0.756) | 2.94<br>(0.300) | 2.06<br>(0.340) | 2.97<br>(0.816) |
|  | r | Early summer | 0.93<br>(0.019) | 0.92<br>(0.026) | 0.76<br>(0.067) | 0.76<br>(0.032) | 0.77<br>(0.037) | 0.64<br>(0.062) | 0.74<br>(0.030) | 0.72<br>(0.053) | 0.37<br>(0.089) |
|  |  | Late summer | 0.91<br>(0.055) | 0.85<br>(0.074) | 0.61<br>(0.068) | 0.67<br>(0.092) | 0.62<br>(0.092) | 0.48<br>(0.100) | 0.65<br>(0.087) | 0.53<br>(0.106) | 0.00<br>(0.156) |

RMSE = root mean squared error; MAE = median absolute error; r = Pearson's correlation coefficient

Microclimate models were run from 1999 to 2019, which covers the majority of the period for which hourly automated meteorological observation data exists for the island. Models were run for each season separately. Across the 20-year time series, we calculated the average seasonal mean temperature (T_mean_) and the average seasonal growing degree days > 5 °C (GDD5), which are both biologically relevant climate variables. Each of these bioclimate variables was calculated for both the AWS and CRA model predictions.

### Microclimate data in species distribution models

To assess the implications of the observed differences between the AWS and CRA driven microclimate models in an applied setting, we model the percentage cover of a widespread and recently threatened endemic plant species on the island, *Azorella macquariensis*, using spatially contiguous microclimate estimates, rather than terrain proxies (c.f. Bricher et al., 2013). This is a critical step for understanding the effects of climate change on this threatened ecosystem because by building cover models for species using microclimate data—rather than terrain proxies—the sensitivity of the system to changes in the microclimate conditions can be explored. Proportional cover data were collected across 90 sites using a stratified random sampling strategy across the landscape of Macquarie Island. Sampling was conducted across a 700 m^2^ area at each site; for full details see Dickson et al. (2019). Here, we added an additional ten sites drawn randomly from grid cells on the coastal fringes of the island (<50 m asl). *A. macquariensis* is extremely rare below this elevation (where it is outcompeted by other species) and, therefore, the proportion of cover at these sites was set to zero.

Using these data, the proportion of *A. macquariensis* cover was modelled across the entire island as a function of several environmental predictors, with the variables selected based on biological knowledge of the species and insights gained from previous analyses (Dickson et al., 2019, 2020). Latitude and longitude were included to capture spatial variation in soil composition and structure known to affect species distributions but for which no contiguous datasets exist (Adamson, Selkirk, & Seppelt, 1993; B. R. Wilson et al., 2019). GDD5 was selected because plant cover is known to be affected by temperature extremes, including the number of growing days in the short growing season of the region, and we also included a quadratic term GDD5^2^ because visualisation of the raw data suggested an apparent nonlinear association between cover and GDD5. Bayesian beta regression models (link = logit; link phi = identity) were fit using the *rstanarm* package in R (Goodrich, Gabry, Ali, & Brilleman, 2018). The beta distribution is a continuous probability distribution defined on the interval (0, 1) and, therefore, response variables must be transformed if they include exact values of 0 or 1. We used the transformation (*y* ×(*n*−1)+0.5/*n*, where *y* is the response variable and *n* is the sample size, to rescale the response variables onto the (0, 1) interval (Cribari-Neto & Zeileis, 2009). Leave-one-out cross-validation (n = 1000) was used to evaluate the model predictive performance. Models were built using both the AWS and CRA driven microclimate datasets and the median posterior predictions for percentage cover were compared by calculating the pair-wise Pearson’s correlation coefficient.

## RESULTS

### Microclimate Model Evaluation

Models driven by AWS data predicted the seasonal T_mean_ across the island with a RMSE ≤ 0.6 °C and MAE ≤ 0.5 °C in both the early and late summer (Table 2). Predictions errors were consistently lower in the late summer. RMSEs for seasonal T_min_ and T_max_ were all <1°C for models driven by the AWS data (RMSE for T_min_ and T_max_ in the late summer ≤0.3 °C and ≤0.8 °C, respectively). Predictions from the CRA model had higher prediction errors than those from the AWS model in the early summer, but had comparable error rates in the late summer (i.e. RMSE for late summer T_mean_ = 0.27 vs. 0.45; Table 2). The daily RMSE for T_mean_ indicates a mean daily error rate for predictions from the AWS models of c. 0.9–1.2 °C (depending on the season) and higher error rates for the CRA model daily predictions (c. 1.2– 1.7 °C). Prediction errors were comparable for daily T_min_ and T_max_. The correlation between observed and predicted values for daily T_min_ and T_mean_ were consistently high for AWS models (r = 0.85–0.93), but lower for predictions of daily T_max_ (r = 0.61–0.76) from these same models, and for T_min,_ T_mean_, and T_max_ from CRA models (r = 0.48–0.77). Microclimate models consistently added value over the macroclimate temperature time-series (NCEP2 derived), which had much higher RMSE rates (T_mean_ > 1.4 °C) and lower temporal correlations for daily temperatures (r = 0.00–0.74) when benchmarked against the in situ observation (Obs) data (Table 2).

Figure 1 shows the predicted and observed temperature time-series, along with the macroclimate temperature time-series, for three randomly selected sites as an illustration of the model predictions. In the early summer, there is no significant difference in the mean seasonal diurnal range predicted across the 62 sites using either the AWS or CRA models (paired t-test; mean dif. = 0.02, t = 1.23, df = 61, p = 0.223), but there is a significant bias in the CRA predictions towards the warmer macroclimate conditions (mean dif. of T_mean_ = 0.72, t = 298.4, df = 61, p ≪ 0.001). This warm bias can clearly be seen in Figure 1. The predictions from the two models in the late summer show a smaller but still statistically significant warm bias in the CRA model predictions (mean dif. of T_mean_ = 0.39, t = 166.4, df = 61, p ≪ 0.001) and a significantly larger diurnal range for the AWS predictions (mean dif. = 0.75, t = 40.42, df = 61, p ≪ 0.001). For all models, the microclimate predictions track the macroclimate temperature variation broadly, but the microclimate models are able to predict more accurately the larger diurnal range experienced on the island. Predictions across all periods do not capture some of the spikes in the daily observed temperature; however, it is possible that some of these spikes are measurement errors in the sensors and not entirely attributable to model errors.

**Figure 1.**
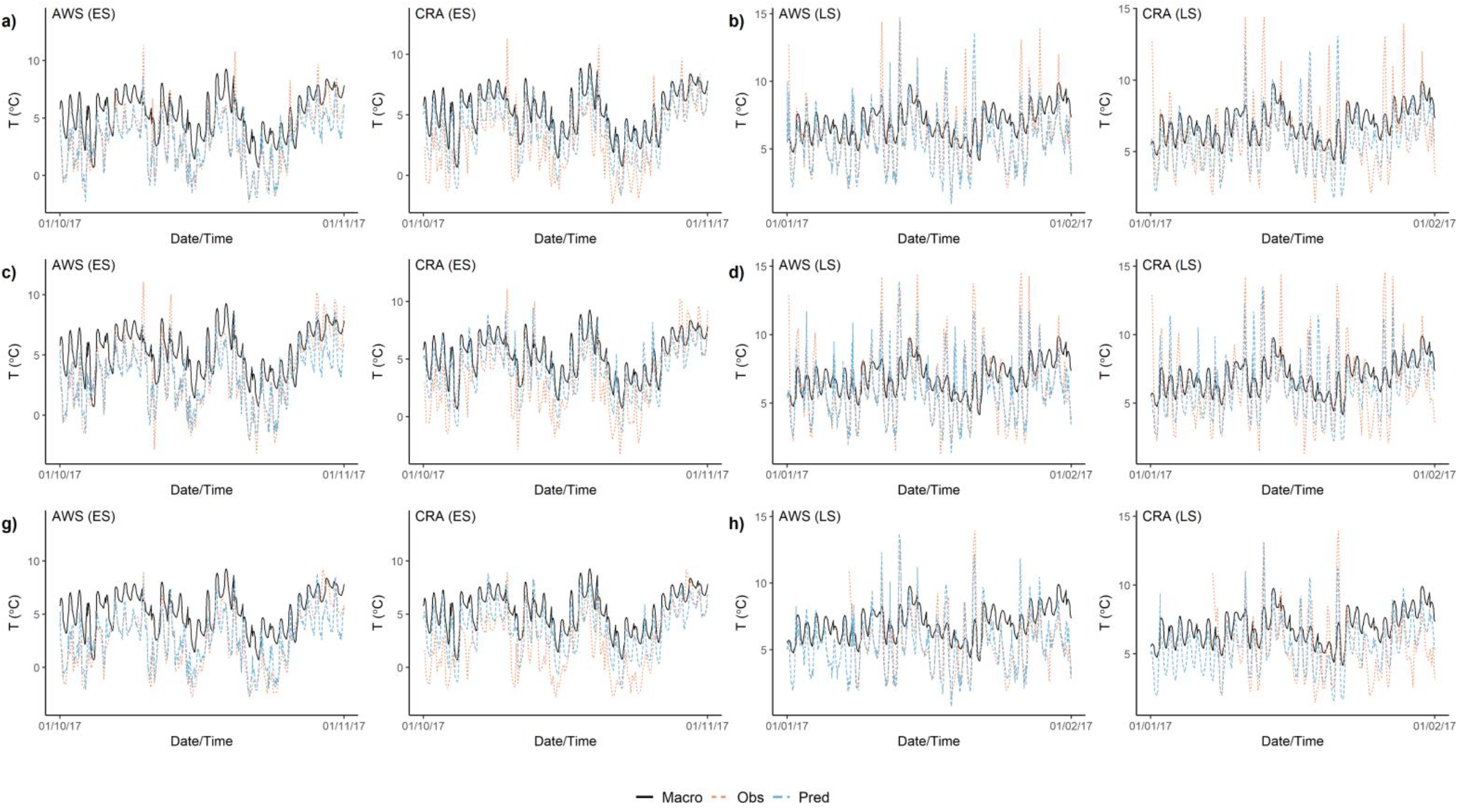
Comparison of fine-scale temperature predictions (Pred) with *in situ* microclimate data logger time-series (Obs) and macroclimate (NCEP2) air temperature (Macro) at three randomly selected sites in the north (a, b), centre (c, d), and south (e, f) of the island for one month in each of the early (ES; Oct-Dec) and late (LS; Jan-Mar) summer periods. Microclimate models were parameterised using either automated weather station (AWS) or climate reanalysis (CRA) data.

Spatial patterns in prediction errors varied between seasons and between models (Fig. 2). Seasonal T_min_ was generally over predicted (Fig. 2a-d), although prediction errors from the AWS model in the late summer showed a slightly more random distribution around zero (Fig. 2c). There was no strong pattern in the size of the T_min_ error across the island for either model or season. The prediction errors for seasonal T_mean_ are generally small and distributed around zero for predictions from the AWS model in the early summer and CRA model in the late summer (Fig. 2e,h), but show consistent directional errors for the other two T_mean_ spatial predictions (Fig. 2f,g). The prediction errors for seasonal T_max_ follow a similar pattern to those of T_mean_, but tend to be larger and to have greater spatial variation in the size of the prediction errors (Fig. 2i,l). These characteristics are observable in a comparison of the 20-year averaged bioclimate variables, which shows that the CRA model consistently predicts higher T_mean_ and GDD5 than the AWS model (Fig. 3). Even where predictions are biased, the spatial correlation between the AWS and CRA derived bioclimate variables was high (r ≥ 0.95), suggesting that both models predict similar spatial temperature patterns.

**Figure 2.**
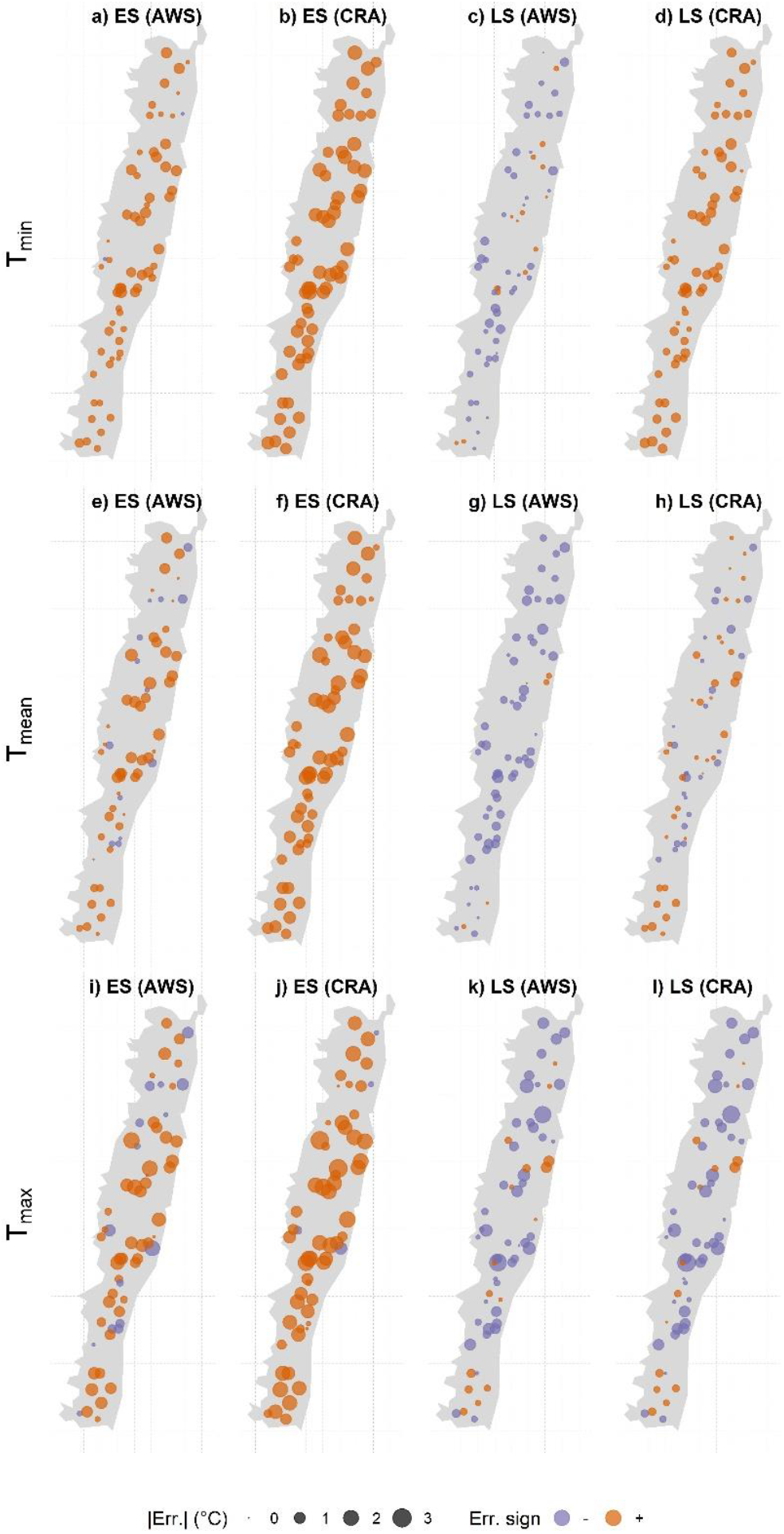
The spatial variation in prediction error for seasonal (early summer [ES] and late summer [LS]) T_min_, T_mean_, and T_max_ across the 62 monitoring sites for microclimate models driven by automated weather station (AWS) or climate reanalysis (CRA) data. (-) indicates an underestimate and (+) indicates and overestimate of the seasonal temperature quantile. Errors are measured in °C (and on a continuous scale).

**Figure 3.**
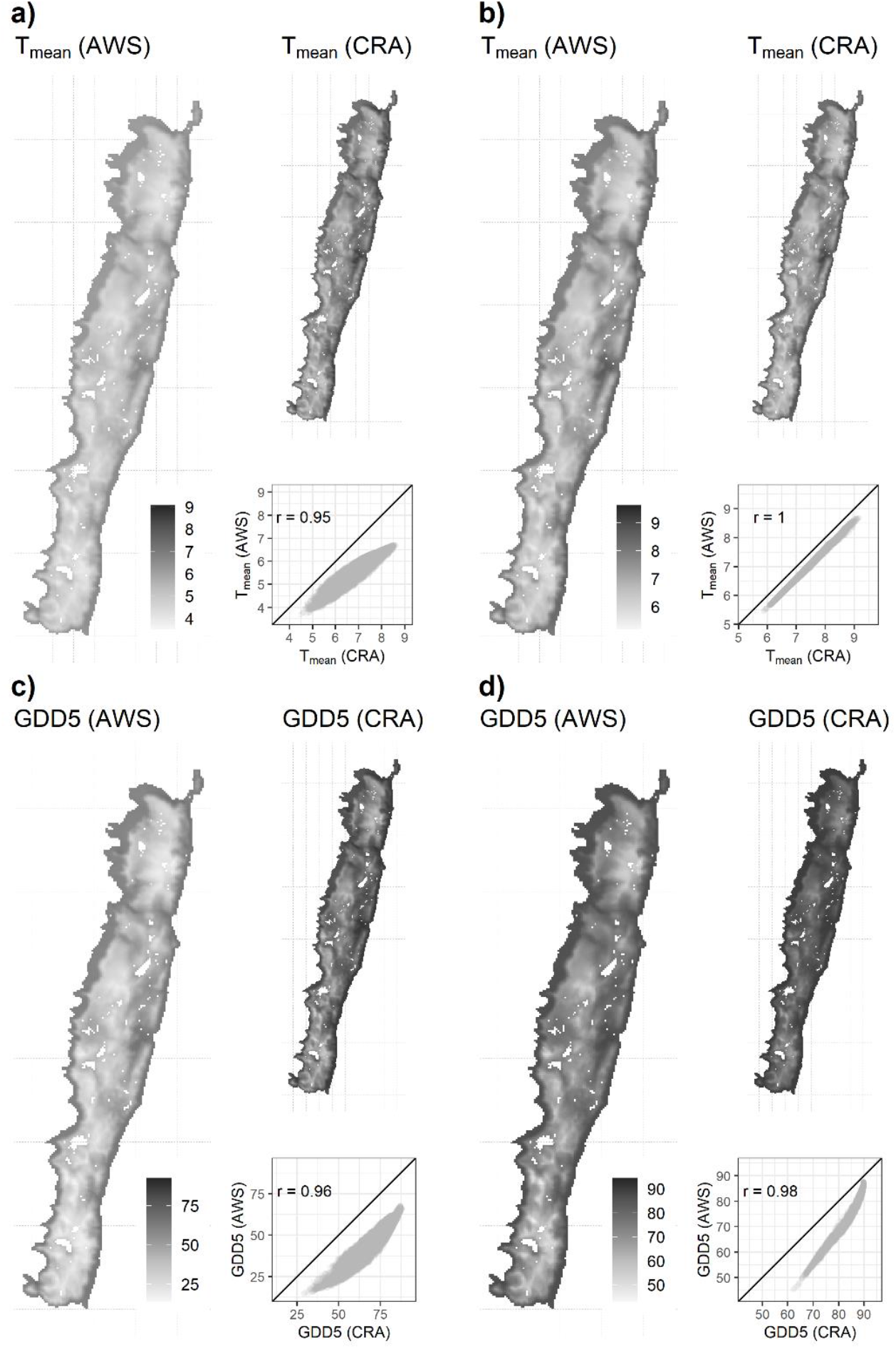
Comparison of bioclimate variables estimated from the microclimate models driven using either automated weather station (AWS) or climate reanalysis (CRA) data, calculated from 20–year (1999–2019) hourly time-series of predicted temperatures at 100 m × 100 m resolution for the (a, c) early summer (ES) and (b, d) late summer periods. T_mean_ = seasonal mean temperature (°C) and GDD5 = average number of seasonal growing degree days >5 °C (number of days). r = Pearson’s correlation coefficient.

### Species distribution modelling

The leave-one-out RMSE for percentage cover predictions using the bioclimate variables derived from the AWS microclimate model had a median RMSE (interdecile range) = 0.11 (0.033, 0.327). This same model fit using the CRA microclimate variables had a median RMSE = 0.13 (0.043, 0.339). The Pearson’s correlation coefficient for the median posterior percentage cover predictions using the two different microclimate datasets was r = 0.97, indicating a very high correlation between predictions from the two models (Fig. 4). Both models predict the locations of major *A. macquariensis* carpets in the south of the island (i.e. around Mt Haswell, Mt Stibbs and Mt Kidson) where three *Azorella* special management areas are delineated to protect high conservation value populations (e.g. high density). Importantly, despite plant cover on the ground being highly variable, other areas of the island with either high density cushion carpets or iconic terraces were also identified by the cover model. While both microclimate models produced highly representative distributions for most populations of *A. macquariensis*, the AWS derived bioclimate variables suggest wider variation in cover in the south and more gradual spatial variation between neighbouring areas.

**Figure 4.**
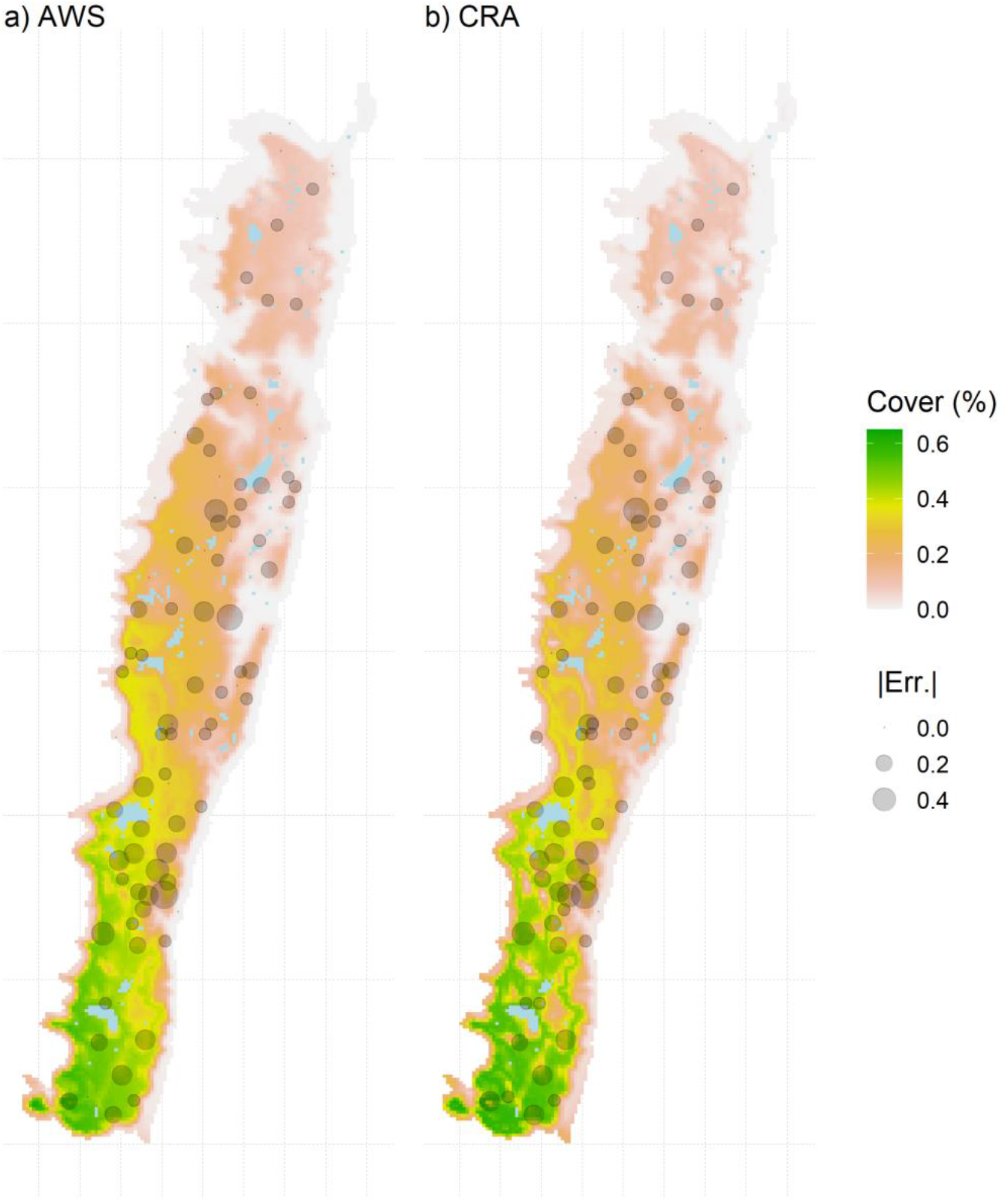
The median posterior predicted cover (%) for the threatened foundation plant species *Azorella macquariensis* made using a beta regression model that included fine-scale bioclimate predictor variables calculated for the early summer period. Bioclimate variables were calculated from fine-scale temperature predictions made using models driven using either AWS (a) or CRA (b) data. The Pearson’s correlation coefficient (r) between cover predictions = 0.97. The size of the points indicate the absolute value of the prediction error between the observed and predicted cover as an indication of model predictive performance (RMSE [obs vs. pred]: a) 0.11 and b) 0.13).

## DISCUSSION

Here, we have evaluated the use of a hybrid physically- and empirically-based modelling approach for predicting daily and seasonal fine-scale climate variation across a remote climate-change threatened polar ecosystem at ecologically relevant spatial scales. When parameterised using *in situ* meteorological observations from the island’s automated weather station and tested against *in situ* microclimate data loggers, model performance was comparable to statistics from other regions (Kearney et al., 2020; Maclean et al., 2017). CRA models, driven using only NCEP2 reanalysis data, had a slight warm bias in the early summer period, but otherwise were able to capture the characteristics of the island’s microclimate variation in space and time. Both models were able to add value (i.e. reduce prediction errors benchmarked against *in situ* observations) to the macroclimate temperature time-series for the island. The application of these data to the challenge of predicting the distribution of a threatened foundation species, *A. macquariensis*, showed that predictions varied little with the source of bioclimate covariates (i.e. AWS or CRA). This suggests that where *in situ* meteorological observations are not available, microclimate models driven only by climate reanalysis data are able to capture the spatial variation in fine-scale climate conditions. These results show that spatially and temporally explicit microclimates may be accurately predicted across remote and exposed landmasses, which paves the way for novel and innovative research on the vulnerabilities and conservation opportunities in these challenging and threatened environments.

The models accurately predict the day-to-day variation in temperature, although there is still error in these predictions and apparent bias in some of the CRA model predictions. The latter is not unexpected as predictions that are unconstrained by *in situ* data are often biased in some respect, and climate models typically go through a bias-correction process prior to use (e.g. Navarro-Racines, Tarapues, Thornton, Jarvis, & Ramirez-Villegas, 2020). Some of this error and bias will be caused by differences between the realised macro-, meso- and micro-scale conditions that influence climate variations across a landscape and those captured by the data used to parameterise the models. For example, here we used cloud cover estimates from the NCEP2 reanalysis data, which have a course spatial (c. 200 × 200 km) and temporal (6 h) resolution and, thus, may underestimate variation in cloud cover, especially over a small landmass situated in a vast ocean. Thus, the reanalysis data may not be entirely representative of the island specific climatic conditions given the dominance of sea in the region (Kearney et al., 2020). Similarly, the summary of orographic information below the grain of the model (here, 100 × 100 m) will introduce variation into the model predictions. The comparison of areal estimates to point estimates (i.e. iButton) within a landscape with even moderately complex orography is likely to increase estimates of prediction error. There can also be considerable error in iButton temperature measurements, depending on the conditions, iButton model, and shielding method employed (Terando, Youngsteadt, Meineke, & Prado, 2017). However, the microclimate model predictions closely match the *in situ* measurements (Fig. 1) and this convergence on the observation data from an independent model suggests that the *in situ* measurements were not strongly afflicted with measurement error.

Despite these uncertainties, these results demonstrate that the hybrid physically- and empirically-based modelling approach, and in particular the AWS driven model, do provide accurate predictions of the seasonal variation (Fig. 1) and seasonal averages (Table 2). The seasonal RMSE for the AWS model compares favourably with predictions of spatially contiguous seasonal climate grids produced with a correlative approach, which ranged from 0.5 to 3.4 °C depending on the quantile being evaluated (Ashcroft, Chisholm, & French, 2009). The difference here is that the model does not require the deployment of hundreds of temperature data loggers for the calibration and model fitting, only data from a single observation station. Also, because the model is not reliant on *in situ* data logger data for calibration we can predict microclimates for time periods when *in situ* microclimate sensor data are not available (i.e. 19 of the 20 years in the time-series predicted here). The RMSE produced here were smaller than RMSE of satellite remote sensing (c. 1 km grain) tested on Marion Island in the Southern Ocean (Leihy, Duffy, Nortje, et al., 2018), although the difference in grain size (100 m vs 1 km) makes a direct comparison difficult. Thus, the hybrid physically- and empirically-based modelling approach appears to offer performance and logistical advantages over other commonly used methods used to obtain microclimate data.

These performance and logistical advantages are important even beyond the isolated polar regions because microclimate data are needed to underpin fundamental and applied ecological research across the breadth of terrestrial ecosystems and species groups (e.g. Jucker et al., 2020; Nowakowski, Frishkoff, Agha, Todd, & Scheffers, 2018). In particular, conservation strategies centred around identifying potential microrefugia have received much interest, especially for species that cannot track shifting climate niches or adapt *in situ* to changing climate conditions (Ashcroft, 2010; Wilson, Walters, Mayle, Lingner, & Vibrans, 2019). Importantly, by understanding the role of microclimates in structuring biodiversity— e.g. through effects on species physiology, demography, and behaviour—it may also be possible to modify landscapes to increase their climate change resilience (Jucker et al., 2020; Shoo et al., 2011). These research questions are most pressing given the stress that climate change is placing already placing on biodiversity (Descamps et al., 2017; Stephens et al., 2016). Thus, the capability to accurately predict spatially and temporally contiguous microclimates at scales relevant to a focal system or species is the most significant outcome of these technical advances because microclimate datasets can now be rapidly produced for locations and time period that correspond with existing ecological dataset where contemporaneous *in situ* microclimate measurements are not available (Lembrechts & Lenoir, 2020).

In this study we have demonstrated the ability of physically- and empirically-based microclimate models to predict ecologically meaningful microclimate conditions across a remote, exposed, and climate change-vulnerable island ecosystem, particularly when *in situ* meteorological data are available to parameterise the models. This paves the way for novel ecological and biogeographic studies on the role of microclimate in determining biodiversity patterns and trends. Projections of future changes in microclimate conditions are also possible (i.e. Maclean, 2020) and these will be invaluable for understanding the plausible range of changes in microclimate conditions across the region and for predicting the threats these changes pose to biodiversity (e.g. Lembrechts, Nijs, & Lenoir, 2018). These advances will help provide valuable information for delivering robust and climate change-resilient conservation planning that accounts for the critically important microclimate dimension.

## ACKNOWLEDGEMENTS

We thank Aleks Terauds, Ben Raymond and Justine Shaw for discussion and logistical advice. Kate Kiefer provided logistical and field support. John Burgess, Rowena Hannaford and the 69^th^ and 70^th^ Macquarie Island Australian Antarctic Program teams helped establish and retrieve the data-loggers. This research was funded by the Australian Antarctic Division and the Australian Government under an Australian Antarctic Science Program Grant #4312, and support to CRD from an Australian Government Research Training Program (RTP) Scholarship. iButton microclimate data is archived at https://data.aad.gov.au/metadata/records/AAS_4312_MI_microclimate_data_Dec16Dec17_6 2sites

## REFERENCES

Adams, N. (2009). Climate trends at Macquarie Island and expectations of future climate change in the sub-Antarctic. Papers and Proceedings of the Royal Society of Tasmania, 143(1–2), 1–8. https:://doi.org/10.26749/rstpp.143.1.1

Adamson, D. A., Selkirk, J. M., & Seppelt, R. D. (1993). Serpentinite, harzburgite, and vegetation on subantarctic Macquarie Island. Arctic and Alpine Research, 25(3), 216– 219.

Ashcroft, M. B. (2010). Identifying refugia from climate change. Journal of Biogeography, 37(8), 1407–1413. https:://doi.org/10.1111/j.1365-2699.2010.02300.x

Ashcroft, M. B., Chisholm, L. A., & French, K. O. (2009). Climate change at the landscape scale: Predicting fine-grained spatial heterogeneity in warming and potential refugia for vegetation. Global Change Biology, 15(3), 656–667. https:://doi.org/10.1111/j.1365-2486.2008.01762.x

Bergstrom, D. M., Bricher, P. K., Raymond, B., Terauds, A., Doley, D., McGeoch, M. A., … Ball, M. C. (2015). Rapid collapse of a sub-Antarctic alpine ecosystem: The role of climate and pathogens. Journal of Applied Ecology, 52(3), 774–783. https:://doi.org/10.1111/1365-2664.12436

Bergstrom, D. M., & Chown, S. L. (1999). Life at the front: History, ecology and change on southern ocean islands. Trends in Ecology and Evolution, 14(12), 472–477. https:://doi.org/10.1016/S0169-5347(99)01688-2

Bricher, P. K., Lucieer, A., Shaw, J., Terauds, A., & Bergstrom, D. M. (2013). Mapping sub-antarctic cushion plants using random forests to combine very high resolution satellite imagery and terrain modelling. PLoS ONE, 8(8), 1–16. https:://doi.org/10.1371/journal.pone.0072093

Bütikofer, L., Anderson, K., Bebber, D. P., Bennie, J. J., Early, R. I., & Maclean, I. M. D. (2020). The problem of scale in predicting biological responses to climate. Global Change Biology, (July), 6657–6666. https:://doi.org/10.1111/gcb.15358

Cahoon, S. M. P., Sullivan, P. F., Shaver, G. R., Welker, J. M., & Post, E. (2012). Interactions among shrub cover and the soil microclimate may determine future Arctic carbon budgets. Ecology Letters, 15(12), 1415–1422. https:://doi.org/10.1111/j.1461-0248.2012.01865.x

Chown, S. L., Clarke, A., Fraser, C. I., Cary, S. C., Moon, K. L., & McGeoch, M. A. (2015). The changing form of Antarctic biodiversity. Nature, 522(7557), 431–438. https:://doi.org/10.1038/nature14505

Clem, K. R., Fogt, R. L., Turner, J., Lintner, B. R., Marshall, G. J., Miller, J. R., & Renwick, J. A. (2020). Record warming at the South Pole during the past three decades. Nature Climate Change. https:://doi.org/10.1038/s41558-020-0815-z

Cribari-Neto, F., & Zeileis, A. (2009). Beta regression in R. Journal of Statistical Software, 34(2), 1–24. https:://doi.org/10.18637/jss.v034.i02

Descamps, S., Aars, J., Fuglei, E., Kovacs, K. M., Lydersen, C., Pavlova, O., … Strøm, H. (2017). Climate change impacts on wildlife in a High Arctic archipelago – Svalbard, Norway. Global Change Biology, 23(2), 490–502. https:://doi.org/10.1111/gcb.13381

Dickson, C. R., Baker, D. J., Bergstrom, D. M., Bricher, P. K., Brookes, R. H., Raymond, B., … McGeoch, M. A. (2019). Spatial variation in the ongoing and widespread decline of a keystone plant species. Austral Ecology, 44, 891–905. https:://doi.org/10.1111/aec.12758

Dickson, C. R., Baker, D. J., Bergstrom, D. M., Brookes, R. H., Whinam, J., & McGeoch, M. A. (2020). Widespread dieback in a foundation species on a sub-Antarctic World Heritage Island: Fine-scale patterns and likely drivers. Austral Ecology. https://doi.org/10.1111/aec.12958

Dobrowski, S. Z. (2011). A climatic basis for microrefugia: The influence of terrain on climate. Global Change Biology, 17(2), 1022–1035. https:://doi.org/10.1111/j.1365-2486.2010.02263.x

Duffy, G. A., Coetzee, B. W. T., Latombe, G., Akerman, A. H., McGeoch, M. A., & Chown, S. L. (2017). Barriers to globally invasive species are weakening across the Antarctic. Diversity and Distributions, 23(9), 982–996. https:://doi.org/10.1111/ddi.12593

Goodrich, B., Gabry, J., Ali, I., & Brilleman, S. (2018). rstanarm: Bayesian applied regression modelling via Stan. R Package Version 2.17. 4.

Hay, J. E., & McKay, D. C. (1985). Estimating solar irradiance on inclined surfaces: A review and assessment of methodologies. International Journal of Solar Energy, 3(4–5), 203–240. https:://doi.org/10.1080/01425918508914395

Hilde, C. H., Christophe, P., Høyvik Hilde, C., Pélabon, C., Guéry, L., Gabrielsen, G. W., & Descamps, S. (2016). Mind the wind: Microclimate effects on incubation effort of an arctic seabird. Ecology and Evolution, 6(7), 1914–1921. https:://doi.org/10.1002/ece3.1988

Jucker, T., Jackson, T. D., Zellweger, F., Swinfield, T., Gregory, N., Williamson, J., … Coomes, D. A. (2020). A research agenda for microclimate ecology in human-modified tropical forests. Frontiers in Forests and Global Change, 2, 1–11. https:://doi.org/10.3389/ffgc.2019.00092

Kanamitsu, M., Ebisuzaki, W., Woollen, J., Yang, S.-K., Hnilo, J. J., Fiorino, M., & Potter, G. L. (2002). NCEP-DOE AMIP-II Renalalysys (R-2). Bulletin of the American Meteorological Society, 83(11), 1631–1643. https:://doi.org/10.1175/BAMS-83-11

Kankaanpää, T., Abrego, N., Vesterinen, E., & Roslin, T. (2020). Microclimate structures communities, predation and herbivory in the High Arctic. Journal of Animal Ecology, n/a(n/a). https:://doi.org/10.1111/1365-2656.13415

Kearney, M. R., Gillingham, P. K., Bramer, I., Duffy, J. P., & Maclean, I. M. D. (2020). A method for computing hourly, historical, terrain-corrected microclimate anywhere on earth. Methods in Ecology and Evolution, 11(1), 38–43. https:://doi.org/10.1111/2041-210X.13330

Kearney, M. R., & Porter, W. P. (2017). NicheMapR – an R package for biophysical modelling: the microclimate model. Ecography, 40(5), 664–674. https:://doi.org/10.1111/ecog.02360

Kemp, M. U., Emiel van Loon, E., Shamoun-Baranes, J., & Bouten, W. (2012). RNCEP: Global weather and climate data at your fingertips. Methods in Ecology and Evolution, 3(1), 65–70. https:://doi.org/10.1111/j.2041-210X.2011.00138.x

Leihy, R. I., Duffy, G. A., & Chown, S. L. (2018). Species richness and turnover among indigenous and introduced plants and insects of the Southern Ocean Islands. Ecosphere, 9(7), e02358. https:://doi.org/10.1002/ecs2.2358

Leihy, R. I., Duffy, G. A., Nortje, E., & Chown, S. L. (2018). Data descriptor: High resolution temperature data for ecological research and management on the Southern Ocean Islands. Scientific Data, 5, 1–13. https:://doi.org/10.1038/sdata.2018.177

Lembrechts, J. J., & Lenoir, J. (2020). Microclimatic conditions anywhere at any time! Global Change Biology, 26(2), 337–339. https:://doi.org/10.1111/gcb.14942

Lembrechts, J. J., Nijs, I., & Lenoir, J. (2018). Incorporating microclimate into species distribution models. Ecography, 42(7), 1267–1279. https:://doi.org/10.1111/ecog.03947

Maclean, I. M. D. (2020). Predicting future climate at high spatial and temporal resolution. Global Change Biology, 26(2), 1003–1011. https:://doi.org/10.1111/gcb.14876

Maclean, I. M. D., Hopkins, J. J., Bennie, J., Lawson, C. R., & Wilson, R. J. (2015). Microclimates buffer the responses of plant communities to climate change. Global Ecology and Biogeography, 24(11), 1340–1350. https:://doi.org/10.1111/geb.12359

Maclean, I. M. D., Mosedale, J. R., & Bennie, J. J. (2019). Microclima: An r package for modelling meso- and microclimate. Methods in Ecology and Evolution, 10(2), 280–290. https:://doi.org/10.1111/2041-210X.13093

Maclean, I. M. D., Suggitt, A. J., Wilson, R. J., Duffy, J. P., & Bennie, J. J. (2017). Fine-scale climate change: Modelling spatial variation in biologically meaningful rates of warming. Global Change Biology, 23(1), 256–268. https:://doi.org/10.1111/gcb.13343

McClelland, G. T. W., Altwegg, R., Van Aarde, R. J., Ferreira, S., Burger, A. E., & Chown, S. L. (2018). Climate change leads to increasing population density and impacts of a key island invader. Ecological Applications, 28(1), 212–224. https:://doi.org/10.1002/eap.1642

Navarro-Racines, C., Tarapues, J., Thornton, P., Jarvis, A., & Ramirez-Villegas, J. (2020). High-resolution and bias-corrected CMIP5 projections for climate change impact assessments. Scientific Data, 7(1), 7. https:://doi.org/10.1038/s41597-019-0343-8

Niittynen, P., Heikkinen, R. K., Aalto, J., Guisan, A., Kemppinen, J., & Luoto, M. (2020). Fine-scale tundra vegetation patterns are strongly related to winter thermal conditions. Nature Climate Change, 10(12), 1143–1148. https:://doi.org/10.1038/s41558-020-00916-

Niskanen, A. K. J., Niittynen, P., Aalto, J., Väre, H., & Luoto, M. (2019). Lost at high latitudes: Arctic and endemic plants under threat as climate warms. Diversity and Distributions, 25(5), 809–821. https:://doi.org/10.1111/ddi.12889

Nowakowski, A. J., Frishkoff, L. O., Agha, M., Todd, B. D., & Scheffers, B. R. (2018). Changing thermal landscapes: Merging climate science and landscape ecology through thermal biology. Current Landscape Ecology Reports, 3(4), 57–72. https:://doi.org/10.1007/s40823-018-0034-8

Nyakatya, M. J., & McGeoch, M. A. (2008). Temperature variation across Marion Island associated with a keystone plant species (Azorella selago Hook. (Apiaceae)). Polar Biology, 31(2), 139–151. https:://doi.org/10.1007/s00300-007-0341-8

Selkirk, P. M., Seppelt, R. D., & Selkirk, D. R. (1990). Subantarctic Macquarie Island: environment and biology. New York.: Cambridge Press University Press.

Shoo, L. P., Olson, D. H., Mcmenamin, S. K., Murray, K. A., Van Sluys, M., Donnelly, M. A., … Hero, J. M. (2011). Engineering a future for amphibians under climate change. Journal of Applied Ecology, 48(2), 487–492. https:://doi.org/10.1111/j.1365-2664.2010.01942.x

Stephens, P. A., Mason, L. R., Green, R. E., Gregory, R. D., Sauer, J. R., Alison, J., … Willis, S. G. (2016). Consistent response of bird populations to climate change on two continents. Science, 352(6281), 84–87. https:://doi.org/10.1126/science.aac4858

Storlie, C., Merino-Viteri, A., Phillips, B., VanDerWal, J., Welbergen, J., & Williams, S. (2014). Stepping inside the niche: Microclimate data are critical for accurate assessment of species’ vulnerability to climate change. Biology Letters, 10(9), 6–9. https:://doi.org/10.1098/rsbl.2014.0576

Suggitt, A. J., Wilson, R. J., Isaac, N. J. B. B., Beale, C. M., Auffret, A. G., August, T., … Maclean, I. M. D. D. (2018). Extinction risk from climate change is reduced by microclimatic buffering. Nature Climate Change, 8(August). https:://doi.org/10.1038/s41558-018-0231-9

Terando, A. J., Youngsteadt, E., Meineke, E. K., & Prado, S. G. (2017). Ad hoc instrumentation methods in ecological studies produce highly biased temperature measurements. Ecology and Evolution, 7(23), 9890–9904. https:://doi.org/10.1002/ece3.3499

Turner, J., Barrand, N. E., Bracegirdle, T. J., Convey, P., Hodgson, D. A., Jarvis, M., … Klepikov, A. (2014). Antarctic climate change and the environment: An update. Polar Record, 50(3), 237–259. https:://doi.org/10.1017/S0032247413000296

Wauchope, H. S., Shaw, J. D., Varpe, Ø., Lappo, E. G., Boertmann, D., Lanctot, R. B., & Fuller, R. A. (2017). Rapid climate-driven loss of breeding habitat for Arctic migratory birds. Global Change Biology, 23(3), 1085–1094. https:://doi.org/10.1111/gcb.13404

Wilson, B. R., Wilson, S. C., Sindel, B., Williams, L. K., Hawking, K. L., Shaw, J., … Kristiansen, P. (2019). Soil properties on sub-Antarctic Macquarie Island: Fundamental indicators of ecosystem function and potential change. Catena, 177, 167–179.

Wilson, O. J., Walters, R. J., Mayle, F. E., Lingner, D. V., & Vibrans, A. C. (2019). Cold spot microrefugia hold the key to survival for Brazil’s Critically Endangered Araucaria tree. Global Change Biology, 25(12), 4339–4351. https:://doi.org/10.1111/gcb.14755

